# How delayed treatment benefits and harms would impact the optimal timing of statin initiation for cardiovascular primary prevention

**DOI:** 10.1101/608216

**Authors:** Rodney A. Hayward, Greggory Schell, Jennifer G. Robinson, Jeremy B. Sussman, Mariel S. Lavieri

## Abstract

**Background:** Genetic studies suggest that the relative risk reduction (RRR) of statins may increase over time, potentially resulting in much greater long-term benefit if statins are started before cardiovascular (CV) risk is high.

**Methods:** We used a nationally representative sample of American adults to estimate effects of initiating a statin when 10-year CV risk reaches 5%, 10% or 15%. We examined scenarios in which a statin’s initial RRR (30%) gradually doubles over 10 to 30 years of treatment.

**Results:** Initiating a statin when 10-year CV risk is 5% resulted in a mean of 20.1 years on a statin before age 75 (8 years more than starting when CV risk reaches 10%). If a statin’s RRR doubles over 20 years, starting when CV risk is 5% would save about 5.1 to 6.1 additional QALYs per 1000 additional treatment years than starting when CV risk is 10%. Most of this additional benefit was accrued by those who reach a 5% risk at a younger age. Due to the prolonged treatment period, however, early treatment could also result in net harm *if* the treatment slowly increased a major complication of aging, such as muscular or neurological aging.

**Conclusions:** In a thought experiment exploring the impact of delayed effects, we found that if the relative effectiveness of statin therapy gradually doubles over a 10 to 30 year period, starting a statin when 10-year CV risk is 5% could have much more long-term benefit than starting a statin when CV risk is 10%. Most of the additional benefit occurred in those at elevated age-adjusted CV risk. Unfortunately, given the long duration of treatment, substantial delayed statin harms, if present, could outweigh these potential benefits and result in substantial net harm.

## Introduction

A key principle of preventive medicine is that intervening to prevent early disease from developing is often much more effective than treating disease complications once the disease has developed. In this regard, primary cardiovascular (CV) prevention can be thought of in two ways – preventing hard CV events (heart attacks and strokes) before they happen, or preventing the development of atherosclerosis in the first place. Most guidelines for CV prevention look at benefit over a 5-10 year time horizon, and therefore, focus on short-term CV event prevention.

The 2013 ACC/AHA cholesterol guidelines recommended a 7.5% 10-year hard ASCVD risk as the starting point for strongly considering initiation of a statin.^**1**^ Proponents of earlier use of lipid lowering medications (ie, treating prior to an individual’s overall CV risk being substantially elevated), however, have suggested that treating based on a 10-year time horizon is too short-sighted, and that intervening before substantial atherosclerosis occurs will be much more effective at preventing long-term CV morbidity and mortality.^**2-4**^ In addition to the straight-forward logic of this biological-based argument, substantial quasi-experimental evidence based on Mendelian randomization and high quality cohort studies appear to support the concept that the relative effects of lipid lowering may at least double over time.^**4-6**^

When deciding when to start any medicine, however, it is critically important to consider the tradeoff of potential long-term benefits with potential treatment harms (the nuisance, side effects and potential adverse effects of treatment, often quantified as “disutilities”), which begin immediately and can also accumulate substantially over time. Earlier statin treatment could lead to tens of millions of people being on a daily statin for over 30 years. Despite the vigorous debate on this topic, we can find no rigorous analysis quantifying the magnitude to which earlier or later statin initiation would affect hard patient outcomes if delayed treatment benefits, or harms, are present. To inform current clinical debate and help direct future research on this issue, we used a mathematical model to examine how various temporal delays in a statin achieving its maximum treatment effect and how different levels of treatment harms/disutility, if present, would impact the net benefit of earlier initiation of statins.

## Methods

### Data source and simulated patient population

We derived a large, nationally representative sample of the U.S. population using data from NHANES III (the early 1990s).^**7**^ NHANES III was chosen because it represents a time period in which statin use was rare and blood pressure was treated much less aggressively, allowing for easier estimation of lipid and blood pressure levels in the population if untreated. Using the method of imputation of chained equations,^**8**^ we generated a sample cohort of 100,000 patients. To allow all patients to start off without any CV treatments, for those on antihypertensive medications we estimated the untreated BP by adding the average effectiveness of BP medications to the measured BP in the dataset.^**9**^ We utilized linear regression to forecast each patient’s SBP, HDL and TC change over time, if untreated, up until age 75 (see eAppendix for details). We restricted our analysis to adults over age 30 with a 10-year CV risk below 5%, since this is the lowest CV risk treatment threshold that we evaluated (see *Treatment thresholds*, below).

### Treatment benefits

We examined the likely benefit of treatment with a moderate intensity statin that has an initial relative risk reduction (RRR) of 30%, although a low potency (20% RRR) and high potency (40% RRR) statin were examined in sensitivity analyses.^**10,11**^ We could find no real world data to help guide us on the functional form of the potential increase in a treatment’s RRR over time.^4^ For example, if the estimates from Mendelian randomization studies are correct, such studies have not produced information on whether it would take 10, 20 or even 30 years for treatment to reach its maximum RRR.^**4**^ Therefore, we considered three temporal delay scenarios. First, we examined an instance in which the initial 30% RRR does *<u>not</u>* increase over time (scenario 1). Then, we considered three different temporal delays for the treatment reaching maximum effectiveness; an RRR of 30% for the first 5 years of treatment, followed by a linear incremental doubling to 60% at year 10 of treatment (scenario 2); at year 20 of treatment (scenario 3); and at year 30 of treatment (scenario 4)(see eAppendix for further details).

In our primary analysis, we examined benefits and harms up until age 75 for two reasons. First, there is relatively little evidence on the effectiveness and adverse effects of statins in the general population beyond age 75.^**1**^ Second, evidence suggests that younger individual’s strongly discount life gains that do not occur until they are elderly.^**12,13**^

### Treatment disutilities

We examined a variety of levels of treatment disutility and disutility patterns over time. Available evidence suggests that the short-term (4-5 year) negative effects of being on a statin are limited to the hassle of taking a pill, to mild to moderate side effects that resolve with cessation (mainly myalgias, and possibly difficulty concentrating), and to a very small increase in time to diabetes onset.^**14**^

This suggests that, at least in the short-term, the negative effect of being on treatment is at most low/moderate. Therefore, we examined the impact of a low constant treatment disutility (0.005, 1.8 days of life a year) and a low/moderate disutility (0.01, about 3.6 days a year).^**9,10**^ Since some researchers have argued that younger and healthier patients have considerably higher disutility for taking a medication, we included a fixed level as high as 0.025 in our sensitivity analysis.^15^

We also modeled an instance in which harmful effects of treatments slowly compound over the years. This “slowly increasing treatment harms” scenario was designed to simulate situations in which 10 year negative effects are minimal, but 20 to 30 years of treatment results in 50% to 100% increase in a common complications of aging, such as frailty, diabetes, or degenerative neurological conditions.^**15-19**^ This is meant to account for concerns expressed, even in the short-term, on potential statin effects on muscular health, diabetes, memory loss and peripheral neuropathy. In this scenario, the treatment impacted these conditions minimally initially (disutility = 0.001) but slowly compounds over twenty-five years to a level of 0.05 or 0.1.^**16**^

### Treatment thresholds

We considered treatment strategies that initiated treatment based on a patient’s estimated 10-year hard cardiovascular (CV) risk (heart attacks and strokes) using the Pooled Cohort (ASCVD) risk tool. Although the ACC/AHA guidelines recommend a 7.5% threshold, this is the threshold for initiation of individualized and shared decision-making, not as the point to mandate treatment.^**1**^ Therefore, we modeled thresholds of starting treatment on earlier or later than the 7.5% for initiation of discussion, a 5% versus 10% 10-year hard CV risk.

As a potential approach for identifying the best candidates for early intervention, we examined whether a lower CV risk threshold should be used in those at elevated age-adjusted CV risk (defined using a patient’s estimated 10-year risk at age 50). The 10-y ‘age-50’ CV risk was estimated by entering age 50 into the ASCVD risk equation along with their other risk factors, with the values for blood pressure and lipids adjusted to account for average changes associated with age using multiple regression. This is similar to using lifetime CV risk,^**18**^ but we used age-50 CV risk for three reasons. First, we could not find published full cardiac and stroke equations for estimating lifetime-risk. Second, age-adjusted risk can be more easily estimated using any available risk calculator. Third, this increases emphasis on CV risk due to multiple risk factors and de-emphasizes risk later in life that can be mainly related to age and male sex alone.

### Assessing the benefits of treatment

Our general modeling approach has been described previously,^**9,10,19-21**^ but is briefly described here and in detail in the eAppendix. Each patient begins in the “healthy” state. In each year of follow-up, they could have a hard ASCVD event (heart attack or stroke), which could be fatal or nonfatal. We used the ratio of our treatment’s RRR for non-fatal vs. fatal events that is found for statins. We applied each treatment strategy to each patient for each combination of treatment benefit and disutility function up to age 75. This information was then used to estimate quality-adjusted life-years (QALYs) discounted at 3%, with sensitivity analysis including a discount range from 0% to 5%.

We estimated QALY loss per event using our previously described method.^**9,10,19**^ In brief, we estimated a QALY loss for the year of a CVD event, a smaller QALY loss for each year of life after an event, a rate of fatality per event, and a reduction in life expectancy for each fatal and nonfatal event. Each of these estimates was obtained from published literature and are presented in the eAppendix. Non-cardiovascular mortality (competing risk) was obtained from Centers for Disease Control and Prevention Life Tables.^**7**^

### Analysis

The primary analysis compared marginal effects – the difference in outcomes between initiating a statin at a specific CV risk threshold to waiting to start at the next highest risk threshold. For example, the incremental change in years treated, CV events prevented and QALYs gained for starting the treatment at a 10-year CV risk of 5% was obtained by subtracting the values obtained if treatment was started at 10% from those obtained if the treatment was started at 5%.

Initially, we ran a best-case scenario – statins as the only CV preventive therapy used – to examine the maximum marginal benefit of starting a statin earlier. Next, we examined each statin scenario in association with a simple benefit-based blood pressure treatment approach (two standard blood pressure medications started when CV risk reached 15%). This simple 15% threshold for blood pressure treatment was chosen since it accrues most the population-level benefit of a more complex approach,^**9**^ yet assures that the statin is always started before the blood pressure medications under all three scenarios (the 5%, 10% and 15% thresholds). This allows us to better isolate the impact of different thresholds for statin initiation since BP treatment was always started after the statin. In sensitivity analyses, we examined the impact of using a JNC8 and a full benefit-based tailored treatment approach^**9**^ instead of the simple 2-drug approach.

### Sensitivity analyses

We examined how sensitive our results were to various parameter estimates by varying them across a broad range (see Table 1).

**Table 1.** Base-case estimates and sensitivity analyses range.

| **Parameter of interest** | **Base-case estimate** | **Range assessed by sensitivity analysis** |
| --- | --- | --- |
| 1. Relative Risk reduction (initial/long-term) | 0.3/0.6 | 0.2/0.4 to 0.4/0.8 |
| 2. Treatment disutility | Fixed at 0.005 or 0.01 | Fixed at 0.025 |
|  | Compounds to 0.1 after 25 years | Linearly increase to 0.1 after 25 years |
| 3. Discount rate | 0.03 | 0 to 0.05 |
| 4. Calibration of CV risk prediction tool | No calibration error | 25% under-prediction to 25% over-prediction |
| 5. Effect of non-fatal CV event on increasing CV mortality risk | 20% | 10% to 30% |
| 6. Time horizon for benefit | Up to age 75 | Up to age 85 |

## Results

### Impact of different treatment thresholds on number of years on a statin

Starting a moderate intensity statin when 10-year cardiovascular (CV) risk is 5%, compared to starting at a 10% risk, will on average result in an additional 7.9 years on treatment per person (about 20.1 years total on a statin before age 75) (Table 2). The marginal increase for time on a statin was much less (4.7 years) when comparing starting at a 10-year CV risk of 10%, instead of 15%.

**Table 2.** Impact of different cardiovascular risk thresholds on the total number of treatment years in American adults.

| Start treatment at<br>a 10-y CV risk of:* | Total years treated<br>up until age 75 |
| --- | --- |
| 5% CV risk* | 20.1 |
| 10% CV risk | 12.2 |
| 15% CV risk | 7.5 |

### Benefits of early statin treatment if the relative treatment effect does not increase over time

If we assume that the 30% relative risk reduction (RRR) of a moderate intensity statin does not increase over time, then starting a statin when 10-year CV risk is 5%, compared to 10%, will result in about 1.3 additional QALYs saved per 1000 treatment years if treatment disutility is low, but cause net harm (1.2 QALYs lost per 1000 treatment years) if treatment disutility is low/moderate (.01) (see Table 3). Further, a slowly increasing treatment harm would result in major net harms (4.5 to 11.3 QALYs lost per 1000 treatment years).

**Table 3.** Impact of a statin, with and without a slowly increasing relative risk reduction (RRR), given different thresholds for treatment initiation in American adults (assumes statin therapy is the only primary CV prevention employed).

| Treatment threshold for starting a moderate intensity statin | Marginal increase in QALYs per 1000 treatment-years in people up through age 75* |  |  |  |
| --- | --- | --- | --- | --- |
|  | Treatment Disutility* |  |  |  |
|  | *Constant* |  | *Slow increase to\** |  |
|  | 0.005 | 0.01 | 0.05 | 0.1 |
| **No Delayed Benefit†** |  |  |  |  |
| 5% (vs. 10%) 10-y CV risk | 1.3* | -1.2 | -4.5 | -11.3 |
| 10% (vs. 15%) 10-y CV risk | 3.6 | 1.5 | 1.1 | -2.1 |
| 15% (vs. 20%) 10-y CV risk | 4.7 | 3.4 | 4.1 | 2.0 |
| **10-years until benefit doubles†** |  |  |  |  |
| 5% (vs. 10%) 10-y CV risk | 6.1 | 3.7 | 0.4 | -6.5 |
| 10% (vs. 15%) 10-y CV risk | 8.5 | 6.6 | 6.2 | 3.2 |
| 15% (vs. 20%) 10-y CV risk | 9.8 | 8.1 | 8.8 | 6.8 |
| **20-years until benefit doubles†** |  |  |  |  |
| 5% (vs. 10%) 10-y CV risk | 5.9 | 3.4 | 0.1 | -6.7 |
| 10% (vs. 15%) 10-y CV risk | 7.4 | 5.3 | 5.1 | 1.7 |
| 15% (vs. 20%) 10-y CV risk | 8.1 | 6.8 | 7.1 | 5.4 |
| **30-years until benefit doubles†** |  |  |  |  |
| 5% (vs. 10%) 10-y CV risk | 5.1 | 2.7 | -0.8 | -7.5 |
| 10% (vs. 15%) 10-y CV risk | 6.2 | 4.3 | 4.0 | 0.6 |
| 15% (vs. 20%) 10-y CV risk | 7.4 | 5.4 | 6.1 | 4.1 |

\* In each scenario, the treatment has an initial relative risk reduction (RRR) of 30%. In the “No Delayed Benefit” scenario, RRR remains at 30% indefinitely, whereas in the other scenarios, the RRR slowly increases to 60% over a 10, 20 or 30 year period. The slowly increasing treatment disutilities (to 0.05 and 0.1) represent the type effects that would occur is treatment increased musculo- or neuro-degeneration by 50 to 100%, an effect size that is unlikely to be detectable by post marketing surveillance.
† All results are for marginal effects. For example, in the no delayed benefits scenario, if the treatment disutility is 0.005, then starting a statin when CV risk is 5% will on average yield 1.3 additional QALYs for every 1000 additional treatment years compared to starting when CV risk is 10%. Negative results (in grey) represent the point at which further lowering the threshold for statin initiation results in net harm

### Benefits of early treatment if the treatment effect does increase over time (delayed benefits)

If a standard potency statin’s RRR gradually doubles over time (starts at 30% and reaches a 60% RRR within 10-30 years after initiation) and statin therapy is the only CV preventive treatment used, the marginal benefits of starting at a 5% CV risk, rather than waiting until risk is 10%, increases substantially and is not outweighed by a constant low or low/moderate treatment disutility (2.7 to 6.1 QALYs gained per 1000 treatment years) (Table 3). Even though the benefits of early statin initiation increases substantially under the assumption of delayed benefits, a slowly increasing major treatment harm that reaches a moderate to high treatment disutility after 25 years (0.05 to 0.1) can still result in substantial net harm from early intervention (up to QALYs lost per 1000 treatment years), which was due to the large number of patients who spent more than 20 years on a statin before age 75. However, since waiting until CV risk is 10% greatly reduces the number of patients on statins over 20-30 years, starting at a 10-year CV risk of 10% was always better than waiting until risk reached 15%, even under an assumption of slowly increasing patient harm. The potential benefits of starting when CV risk is 5% go down by about 1 QALY per thousand treatment years if it takes 30 years for a statin to double in effectiveness, rather than it only taking 10 years for a statin to reach full effectiveness (Table 3).

### How higher or lower age-adjusted 10-y CV risk impacts the benefits and risks of earlier statin initiation

Table 4 reports results for the 20-year delayed benefit scenario, but now includes blood pressure treatment once a patient’s CV risk reaches 15% CV risk, and shows results stratified by age-adjusted risk. These results show that those patients with the highest age-adjusted CV risk have the most to gain from starting a statin when CV risk is 5%. For example, for the 25% of the U.S. population with the highest age-adjusted CV risk (made up predominantly of those with multiple risk factors, see eTable 5 in the Appendix), *if* treatment disutility is low or low/moderate (0.005 to 0.01) then roughly 5.8 to 9.7 additional QALYs will be gained per 1000 treatment-years when a statin is started when CV risk is 5%, compared to starting the statin when CV risk is 10%.

**Table 4.** Net benefit of primary CV prevention treatment for American adults if statin’s relative effect doubles over 20 years and blood pressure is treated once patient’s 10-y CV risk is 15%, stratified by age-adjusted risk.*

|  | Treatment threshold<br>for starting a<br>moderate intensity statin | Years on a statin<br>up through<br>age 75 | Net increase in QALYs<br>per 1000 treatment-years |  |  |  |
| --- | --- | --- | --- | --- | --- | --- |
|  |  |  | Treatment Disutility |  |  |  |
|  |  |  | Constant |  | Slow increase<br>to |  |
|  |  |  | 0.005 | 0.01 | 0.05 | 0.1 |
| With Blood pressure treatment |  |  |  |  |  |  |
| All patients | 5% (vs. 10%) 10-y CV risk | 20.1 | 5.3 | 2.9 | -0.5 | -7.4 |
|  | 10% (vs. 15%) 10-y CV risk | 12.2 | 6.2 | 4.3 | 3.8 | 0.6 |
| Top Age-50 CV risk Quartile† | 5% (vs. 10%) 10-y CV risk | 36.7 | 9.7 | 5.8 | -5.8 | -23.4 |
|  | 10% (vs. 15%) 10-y CV risk | 28.4 | 15.3 | 12.3 | 4.0 | -9.2 |
| 2<sup>nd</sup> Age-50 CV risk Quartile | 5% (vs. 10%) 10-y CV risk | 30.2 | 8.1 | 5.0 | -3.7 | -17.2 |
|  | 10% (vs. 15%) 10-y CV risk | 20.7 | 11.9 | 9.6 | 6.1 | -0.5 |
| 3<sup>rd</sup> Age-50 CV risk Quartile | 5% (vs. 10%) 10-y CV risk | 24.5 | 6.2 | 3.6 | -1.3 | -9.9 |
|  | 10% (vs. 15%) 10-y CV risk | 14.3 | 7.5 | 5.6 | 5.4 | 2.5 |
| Bottom Age-50 CV risk Quartile | 5% (vs. 10%) 10-y CV risk | 14.5 | 3.5 | 1.5 | 1.7 | -0.6 |
|  | 10% (vs. 15%) 10-y CV risk | 8.0 | 4.0 | 2.5 | 4.4 | 3.9 |

\* Statin initially reduces CV risk by 30%, but efficacy linearly increases to 60% at 20 years.
† Age-50 CV risk is an age-adjusted risk obtained by putting age 50 into the ASCVD risk calculator regardless of the person’s current age, along with all of their other current risk factors (adjusting for how blood pressure and lipids change with age). It is similar conceptually to risk-stratifying patients based on lifetime risk, but an age-adjusted risk has the advantage of being feasible using any available CV risk calculator. The mean 10-y age-50 CV risk was 18.0% for the “Top Quartile”, 9.8% for the 2<sup>nd</sup> quartile, 6.2% for the 3<sup>rd</sup> quartile, and 3.2% for the bottom quartile. Further details on the age-50 risk quartiles can be found in eTable 5 in the Appendix.

Unfortunately, those with a high age-adjusted CV risk will, on average, spend a very long time on a statin (an average of over 35 years prior to age 75), so they are also at much greater risk of harm if long-term major treatment harms are present. If a major treatment harm slowly compounds and reaches a disutility of 0.05 to 0.1 after 25 years, the highest age-adjusted risk quartile will lose roughly 5.8 to 23.4 QALYs per 1000 treatment-years from starting a statin when CV risk is 5% (Table 4).

### Sensitivity Analyses

Table 5 shows the sensitivity of results across a range of parameter assumptions for the low fixed treatment disutility (0.005) and the delayed high treatment disutility scenarios (reaches 0.1 after 25 years). We found that the former scenario demonstrated net benefit and the latter demonstrated net harm across all parameters assessed, suggesting that our results were robust to model assumptions. However, several factors substantially impacted the magnitude of estimated net benefit and net harm. The factor with the greatest impact on results is how future events are discounted. If the discount rate is 0 (ie, QALYs in the distant future are equal to QALYs in the short-term) then both the net gain of delayed benefits and the net loss of delayed harms both increase substantially (Table 5). In contrast, a discount rate of 5% would make the importance of delayed benefits and delayed harms much less important, making the decision of whether to start treatment at a 5% vs. 10% 10-year CV risk much less important. Extending the time horizon to age 85 had little impact on the net gains of delayed benefits, but would make the risk of delayed harms, if present, much more important (Table 5). This was mainly due to the effects on the low age-adjusted risk group, since in this group most of the delayed harms, if present, would occur between ages 75 and 85 (results not shown).

**Table 5.** Sensitivity analyses for starting a statin when 10-y CV risk is 5% (compared to starting at a 10% CV risk) when a statin’s relative risk reduction (RRR) doubles over a 20-year period.*

| Parameter of interest | Treatment harms fixed (0.005) |  |  | Delayed treatment harm (up to 0.1) |  |  |
| --- | --- | --- | --- | --- | --- | --- |
|  | Basecase | Upper Bound | Lower Bound | Basecase | Upper Bound | Lower Bound |
| 1. Initial RRR 20% to 40%* | 5.35 | 8.62 | 1.92 | -7.38 | -3.93 | -10.48 |
| 2a. Fixed treatment harm 0.025 | 5.35 | 2.93 | -4.07 | -- | -- | -- |
| 2b. Linear increase in delayed treatment harm | -- | -- | -- | -7.38 | -7.38 | -21.13 |
| 3. CV risk tool over- or under-predicts by 20% | 5.35 | 6.48 | 4.06 | -7.38 | -2.12 | -11.78 |
| 4. A non-fatal CV event increases future CV risk by 10% to 30% | 5.35 | 5.73 | 4.96 | -7.38 | -6.87 | -12.16 |
| 5. Time horizon, up to age 75 to up to age 85 | 5.35 | 5.35 | 4.43 | -7.38 | -7.38 | -12.29 |
| 6. Discount Rate, 0% to 5% | 5.35 | 20.24 | 2.04 | -7.38 | -4.07 | -17.95 |

\* A 20% initial RRR simulates results for a low-potency statin and 40% is similar to results for some high potency statins.

## Discussion

Since identifying moderate-sized delayed medication effects is extremely difficult, our study may best be considered a thought experiments, yet demonstrates what is at stake when starting a daily, metabolically active treatment many years before substantive benefits are likely to occur. We found that if the relative treatment effect of a statin gradually doubles over 10 to 30 years and there are no major delayed treatment harms, then starting a statin when 10-y CV risk is 5% will produce much more long-term benefit than waiting until CV risk is 10%. This was true especially for those with a high age-adjusted CV risk (mainly made up of those with multiple risk factors). However, starting intensive statin after CV risk is 10% is a much better choice *if* statins have substantial delayed harms, such as accelerating the rate of a common complication of aging, such as slowly increasing the rate of muscular health or neuro-degeneration.

Statins are one of the most beneficial and commonly used medications ever invented. Although there is a strong consensus that high risk individuals should receive a statin, what the treatment threshold should be for primary prevention is in dispute. There is increasing evidence that overall CV risk should be the main metric for targeting patients for treatment, and recently, guidelines in the U.S. and internationally have proposed recommending a statin when 10-year CV risk reaches about 7.5% to 10%.^**1,22**^ These recommendations have been based mainly on thinking of net benefit over a 10-year horizon. The fact that benefits for those with less than a 10% 10-year CV risk are limited to a modest absolute decrease in CV events, most of which are non-fatal, has made treatment in this group controversial.^**10**^ Some critics have argued that a longer-term horizon is needed, especially given strong pathophysiological, observational and genetic evidence that the relative risk reduction (RRR) from lipid lowering may increase substantially over time.^**2-6**^

The likelihood that delayed benefits or harms exist remain uncertain, but our findings demonstrate that such effects could impact long-term net benefit (or harm) substantially. Therefore, our study might not precisely answer the question regarding the optimal timing of statin initiation, but it can help narrow the debate. If a statin’s RRR does *not* increase over time (ei, you do not believe the genetic studies), our results suggest that there will be only a very small incremental benefit, on average, from starting a statin before 10-year CV risk is 10%. Similar results have been reported by others using a 10-year time horizon.^**10,23**^ Our analyses, however, demonstrate a similarly small benefit would be expected even taking a longer-term view, but also demonstrates just how small this net benefit would be. It would only take a low/moderate treatment harm to result in substantial net harm in this scenario.

Our study is the first, however, to examine how this equation would change if the genetic studies are correct, that the benefits of lipid reductions increase over time.^**2-6**^

- If a statin’s RRR is likely to substantially increase over time *<u>and</u>* you think long-term statin use is unlikely to have substantial harms, then our results suggest that major net benefits will be achieved by starting a statin when CV risk is about 5%, especially for those with a high age-adjusted CV risk.
- In contrast, if a statin’s RRR is likely to substantially increase over time *<u>but</u>* you are very worried about the potential for long-term statin harms, then our results suggest that starting when CV risk is 10%, but not later, is a better approach

Although our paper addresses statin therapy specifically, our results demonstrate more generally how important slowly compounding negative effects can be whenever short-term benefits in clinical trials are limited to non-fatal events, with short-term effects of statin therapy on those with a CV risk below 10% is just one example.^**16,24**^ Clinicians, however, must make decisions based on the information at hand. Even when the range of possibilities is somewhat broad, estimating net benefits under different plausible scenarios can aid shared clinical decision-making and help identify important factors that warrant special attention in future research.

So how likely are statins to have delayed benefits, and how worried should clinicians and patients be about potential delayed statin harms? We expect that opinions on these possibilities will vary widely even between clinical and epidemiological experts. We can only point out that the genetic and epidemiologic evidence suggesting that the RRR of lipid lowering increases substantially over time is very strong,^**2-6**^ and that determining whether a treatment substantially increases long-term harms is a very difficult task. For example, five-years of smoking has little to no impact on a person’s cancer risk, and the delayed harms examined in our study is only about 10% the well-known cancer risk associated with 30 pack-years,^25^ which is too small to be reliably detected given current post-marketing surveillance data sources. This latter point is often under-appreciated, with many feeling that if a treatment has been on the market a long time that major long-term harms would be evident. Observational post-marketing surveillance studies of long-term harms, however, face major methodological difficulties. First, those who experience adverse effects or are less health-conscious are much more likely to discontinue therapy, which can severely confound study results due to a healthy volunteer/adherer effect. For example, high quality observational studies suggested that HRT and beta-carotene were highly beneficial, yet clinical trials demonstrated that these treatments are actually harmful. Major bias was due to healthier people being more likely to be started on and remain on these treatments.^**26,27**^ Such biases can be multiplied several fold when examining outcomes for long-term adherers, ei, those on a treatment for 15-20-years. Further, long-term exposure to statins will be susceptible to potential survival bias (ie, statin treatment can improve survival and therefore those who survive can be sicker on average). Such limitations explain why recent reports that commonly used anti-cholinergic medications increase dementia 2-to 3-fold,^**28-29**^ and that statins increase Parkinson’s disease 2-fold have gone largely, and appropriately, ignored by clinicians.^**30-31**^

Because of this, both post-trial follow-up and cohort studies have difficulty reliably detecting long-term medication adverse effects, unless treatment increases such harms by several fold. Some critics of lowering the statin treatment threshold have raised concerns about whether statins could accelerate muscular de-conditioning, neurodegeneration, and cataracts much more in the long-term than short-term.^**31**^ Further, although current data suggests that statins increase risk of developing diabetes only slightly over a five year period,^**1**^ a slowly compounding effect over 20-25 years could have major negative impact if statins are commonly started in patients in their 40’s and early 50’s. Advocates of early statin intervention, however, can point to some evidence that statins might also have additional long-term benefits, yet critics can point to decades of drug development research that has found that unexpected medication harms are much more common than unexpected medication benefits.

Like all mathematical probability models, our study is limited by the accuracy of highly influential assumptions. Fortunately, other than the key assumption of whether delayed treatment effects exist, our main results were highly robust to large variations in most other model parameters, and most of our assumptions are based on grade B+ to grade A evidence. Our results were most sensitive to the discount rate for future events. If and how much future events should be discounted is a philosophical, not scientific, question, but we do know that patients vary substantially in how much they discount the future.^**12,13,15**^

We fully realize that our results may be frustrating to many clinicians – starting a statin when 10-year CV risk is 5% could be very beneficial or very harmful depending on the presence of uncertain circumstances. The uncertainty surrounding this issue, however, should not distract us from this issue’s clinical importance. An increase or decrease of 5 to 8 QALYs per 1000 treatment years is quite substantial, and lowering the treatment threshold to 5% would result in almost all people being started on a statin before age 65. Further, the group with the most to gain from early treatment, those that reach a 5% CV risk at a younger age, are also those with the most to lose if statin’s have delayed harms. Despite the difficulty of detecting delayed benefits and harms, attention to this issue warrants great attention by researchers given its public health importance.

## Key Points

1. A policy of initiating a statin when 10-y CV risk is 5%, instead of waiting until risk is 10%, would result in an average time spent on a statin of about 20 years prior to age 75 (about 8 years more per person compared to starting at a CV risk of 10%).
2. ***If*** the benefits of a moderate intensity statin slowly double over a 10 to 30 year period ***and*** the long-term negative effects of statin treatment are at most low/moderate, those with a high age-adjusted CV risk will receive substantial net benefit from starting a statin when CV risk is 5% or greater.
3. Due to the prolonged treatment period, however, early treatment can still result in substantial net harm ***if*** treatment slowly increases progression of another common condition of aging, such as frailty or neurodegeneration, over a 25 year period.
4. ***If*** a statin’s relative effect doubles over time, waiting past a 10-y CV risk of 10% results in a substantial net loss of QALYs even in the presence of substantial delayed treatment harms.

## eAppendix

### Mathematical model

We utilized a controlled Markov chain to compute results for the treatment strategies.^**1,2**^ eTable 1 presents the inputs and data sources for the Markov model.

**eTable 1.**
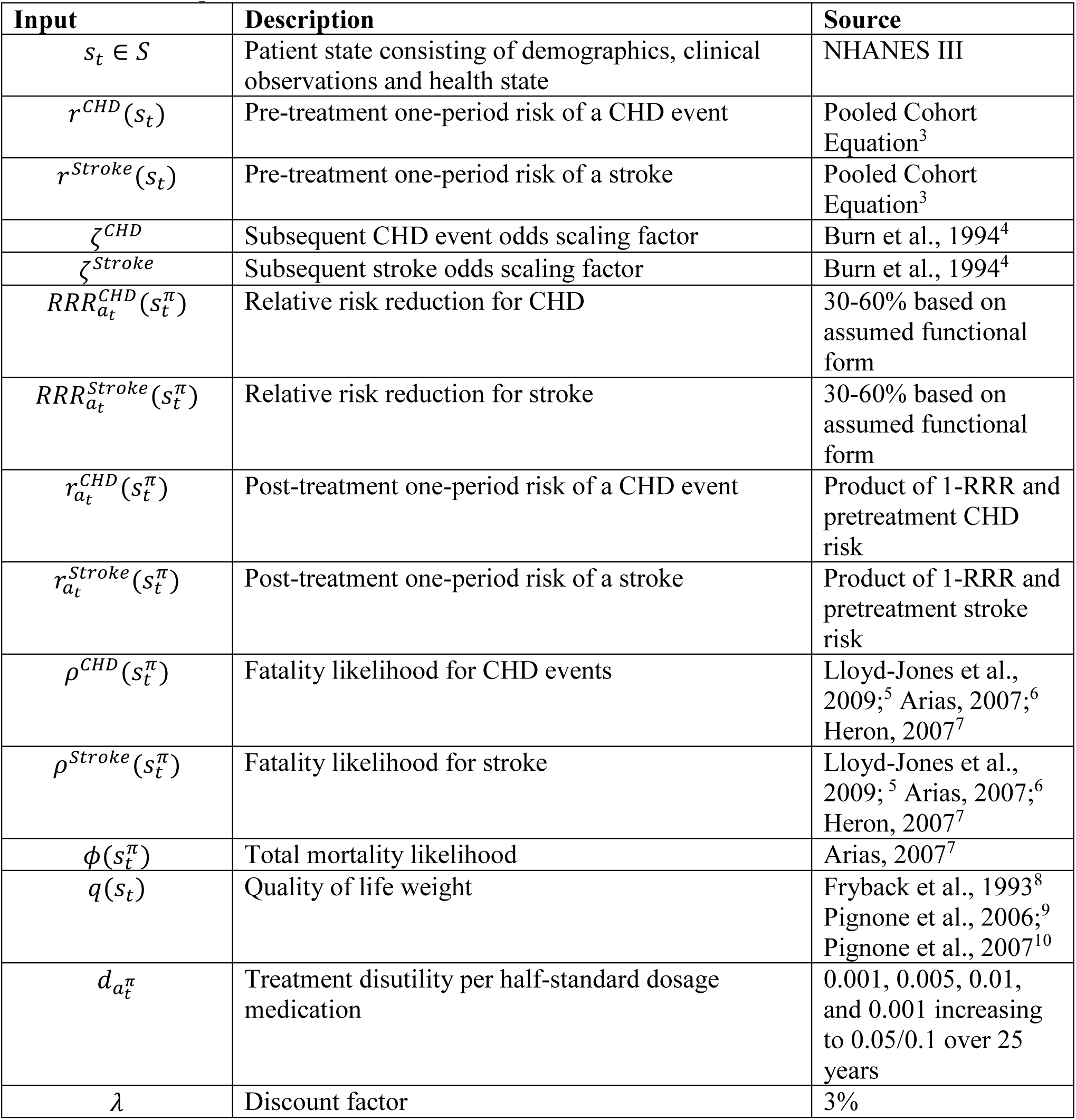
Model Inputs

The Markov chain state^**1,2**^ *st* consisted of demographic information, clinical observations, and the patient’s health state. The demographic information includes the patient’s age, sex, smoking status, and diabetes status. The clinical observations are measurements of the patient’s untreated systolic blood pressure (SBP), high-density lipoprotein (HDL), total cholesterol (TC), and if the patient has left ventricular hypertrophy (LVH) as determined by electrocardiogram. Lastly, there are 10 mutually-exclusive patient health states: (1) healthy (no history of CHD or stroke); (2) history of CHD but no CHD event this period; (3) history of stroke but no stroke this period; (4) history of CHD and stroke but no adverse event this period; (5) survived a CHD event this period; (6) survived a stroke this period; (7) death from a non-CVD related cause; (8) death from CHD event this period; (9) death from stroke this period; and (10) dead.

The clinical observations transitioned deterministically in accordance with the linear regression models developed for each risk factor. The health state transitioned based on the patient’s one-year CV risk and the assumed treatment benefit (once the patient initiated treatment). If the patient has a history of CHD and/or stroke, we multiply the patient’s CHD and stroke odds by a scaling factor *ζ*^*CHD*^ and/or *ζ*^*Stroke*^. We used a scaling factor of 3 for CHD and stroke.^4^ With fatality likelihoods for CHD events *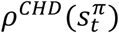* and strokes *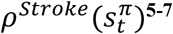* and total mortality Likelihood *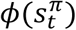*, we computed the probability of transitioning from one health state *h = 1, …, 10* to another health state *h*^*’*^ *= 1, …*, *10*. The fatality likelihoods are presented in eTable 2. We assume only one type of event may happen in a decision period, e.g. a patient cannot have both a CHD event and a stroke. In order to guarantee valid probability distributions, we prioritize death over other outcomes and adjust the health state transition probabilities accordingly.

**eTable 2.** Fatality Likelihoods

| **Age** | **CHD** |  | **Stroke** |  |
| --- | --- | --- | --- | --- |
|  | **Male** | **Female** | **Male** | **Female** |
| 40-44 | 0.12 | 0.26 | 0.03 | 0.04 |
| 45-54 | 0.19 | 0.37 | 0.07 | 0.08 |
| 55-64 | 0.26 | 0.48 | 0.11 | 0.11 |
| 65-74 | 0.35 | 0.64 | 0.16 | 0.16 |
| 75+ | 0.44 | 0.64 | 0.21 | 0.21 |

### Outcome Measures

We computed the number of CV events, discounted QALYs (3% discounting), number of treatment years, and number of people treated for each treatment strategy under each combination of treatment benefit and disutility function. eTable 3 presents the quality of life weights used for each health state.

**eTable 3.** Quality of Life Weights

| Health State | Quality of Life Weight |
| --- | --- |
| Healthy | 1 |
| History of CHD but no event this period | 0.90 |
| History of stroke but no event this period | 0.90 |
| History of CHD and stroke but no event this period | 0.81 |
| Survival of a CHD event this period | 0.88 |
| Survival of a stroke this period | 0.67 |
| Death from any cause | 0 |

At age 75, we computed terminal conditions based upon the ending health state. We assume the terminal condition can be computed as the product of the patient’s expected lifetime,^**6**^ a mortality scaling factor,^**11**^ and the quality of life weighting for the health state. eTable 4 reports the mortality scaling factor and quality of life weights for each terminal health state.

**eTable 4.**
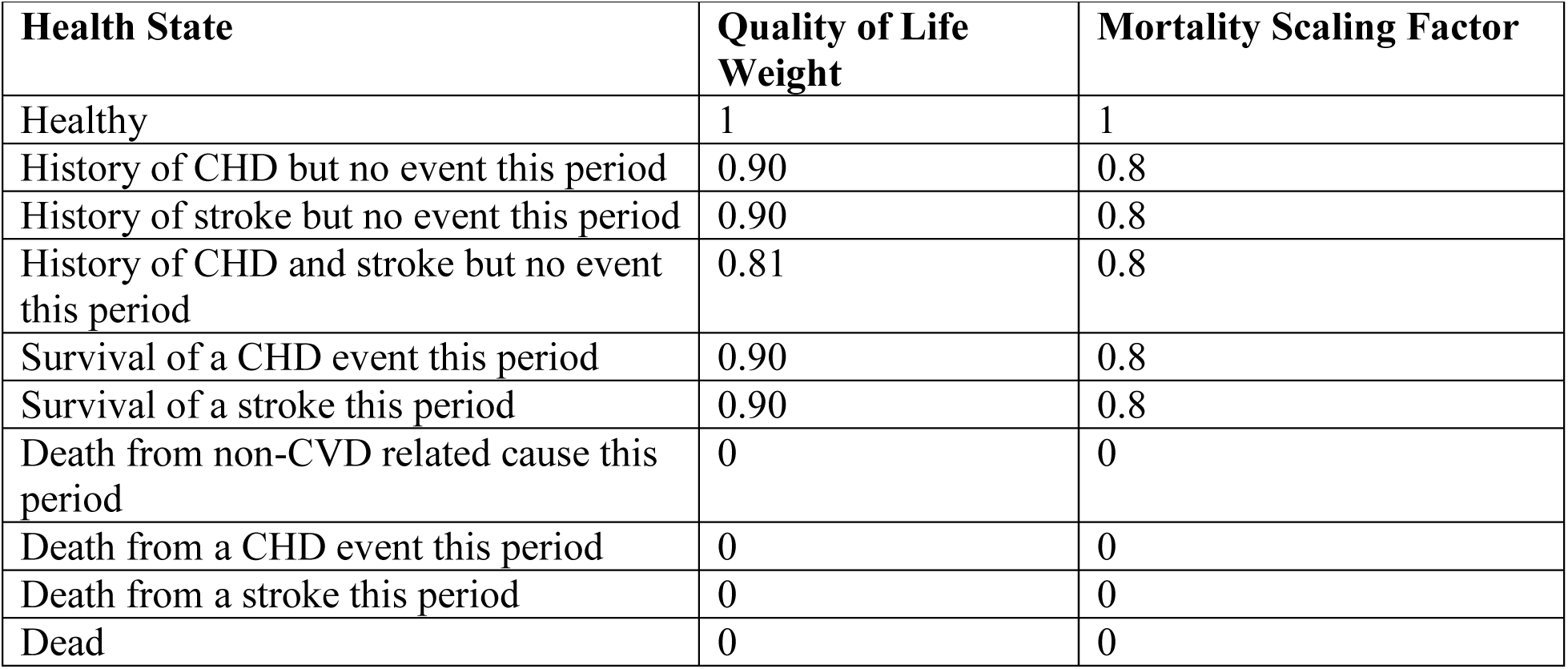
Terminal Condition Inputs

| Health State | Quality of Life Weight | Mortality Scaling Factor |
| --- | --- | --- |
| Healthy | 1 | 1 |
| History of CHD but no event this period | 0.90 | 0.8 |
| History of stroke but no event this period | 0.90 | 0.8 |
| History of CHD and stroke but no event this period | 0.81 | 0.8 |
| Survival of a CHD event this period | 0.90 | 0.8 |
| Survival of a stroke this period | 0.90 | 0.8 |
| Death from non-CVD related cause this period | 0 | 0 |
| Death from a CHD event this period | 0 | 0 |
| Death from a stroke this period | 0 | 0 |
| Dead | 0 | 0 |

**eTable 5.** Attributes of U.S. adults by age-50 CV risk quartiles *

|  | **Top risk quartile** | **2<sup>nd</sup> Risk quartile** | **3<sup>rd</sup> Risk quartile** | **Bottom Risk quartile** |
| --- | --- | --- | --- | --- |
| Female, % | 22.2 | 43.5 | 63.4 | 85.4 |
| Smoker, % | 52.6 | 29.3 | 16.2 | 6.9 |
| HDL < 45mg/dL, % | 74.9 | 48.8 | 28.3 | 9.2 |
| HDL, *mean±sd in mg/dL* | 39.3±10.7 | 46.9±11.4 | 53.1±12.6 | 64.4±16.7 |
| TC, *mean±sd in mg/dL* | 230.4±39.5 | 217.3±40.2 | 209.0±41.6 | 192.6±41.5 |
| Have Diabetes, % | 15.5 | 7.9 | 3.6 | 1.3 |
| SBP > 140mmHg, % | 41.7 | 24.3 | 15.2 | 6.2 |
| SBP, *mean±sd in mmHg* | 137.7±18.2 | 129.4±15.3 | 124.9±14.9 | 117.7±14.1 |
| Estimated 10-y age-50 CV risk,*<br>*mean (range), %* | 18.0<br>(12.3, 77.0) | 9.8<br>(7.7, 12.3) | 6.2<br>(4.7, 7.7) | 3.2<br>(0.4, 4.7) |

\* Age-50 CV risk is an age-adjusted risk obtained by putting age 50 into the ASCVD risk calculator regardless of the person's current age, along with all of their other current risk factors (adjusting for how blood pressure and lipids change with age). It is similar conceptually to risk-stratifying patients based on lifetime risk, but an age-adjusted risk has the advantage of being feasible using any available CV risk calculator. The mean 10-y age-50 CV risk was 18.0% for the "Top Quartile", 9.8% for the 2<sup>nd</sup> quartile, 6.2% for the 3<sup>rd</sup> quartile, and 3.2% for the bottom quartile.

